# Songbird connectome reveals tunneling of migratory neurons in the adult striatum

**DOI:** 10.1101/2025.10.24.684388

**Authors:** N.R. Shvedov, S.J. Castonguay, A. Rother, D.E. Schick, J. Kornfeld, B.B Scott

## Abstract

Immature neurons in the adult brain migrate and integrate into existing circuits, where they contribute to plasticity, learning, and complex behaviors. However, how these cells navigate synapse-rich regions of the adult brain remains poorly understood. While prior studies have examined the molecular mechanisms and functional consequences of adult neurogenesis, few have investigated the physical interactions between migrating neurons and their surrounding environment. Here, we use electron microscopy-based connectomics to examine how migrating neurons interact with mature circuit elements in the adult zebra finch striatum. Immature neurons exhibiting migratory features were observed contacting diverse structures in their microenvironment, including the axons, dendrites, synapses, and somas of mature neurons. Surprisingly, these interactions were structurally complex, often involving pronounced deformations of mature somas and the surrounding neuropil. These deformations appeared as “tunnels” made by the migratory neurons as they displaced mature structures along their path. Together, these findings suggest that migrating neurons may physically reshape the mature circuit to reach their targets, revealing an unexpected degree of structural and functional plasticity in the adult brain.

## Introduction

Adult neurogenesis is believed to play an important role in learning and memory, tissue homeostasis, and regeneration ^1–3^. In many adult vertebrates, this process involves migration, in which immature neurons move from proliferative zones to their integration targets. Mature brain tissue poses several challenges for migrating neurons. Compared with the developing brain, adult brains have increased cell density and diminished extracellular space^4^, and are rich in stable synapses, believed to form the basis of behavior and long-term memory^5,6^. Given these constraints, how migratory neurons can physically navigate through the adult brain environment is not clear.

Songbirds are a valuable model organism for the study of neuron migration in the adult brain ^7,8^. In these species, new neurons integrate into brain regions that control complex learned behaviors, where they establish synapses with mature neurons and respond to sensory stimuli ^9–13^. New neurons are added to at least two subregions within the circuit that control learning and production of song--HVC, a pallial premotor nucleus, and Area X, a region within the striatum^14^.

New neurons that migrate to the striatum are born within a region of the lateral ventricle, corresponding to the mammalian medial ganglionic eminence (MGE)^15,16^. After arrival in the striatum, they differentiate into medium spiny neurons (MSN) and integrate into the local circuit ^17–19^. New neurons have also been reported in the adult mammalian striatum^20–22^, motivating further study of neurogenesis in this region in songbirds. However, a key problem is how the new neurons interact with mature circuit structures in the striatum, such as axons, dendrites, and neuronal somas.

Here, we leverage the ultrastructural resolution and dense labeling of electron microscopy (EM) connectomics to investigate the microenvironment of migratory neurons in the adult brain. We focused on the first fully reconstructed zebra finch connectome, collected from the song nucleus Area X in the striatum^23,24^. Within this dataset, we identified a population of migratory neurons using a supervised machine-learning classifier based on morphological features derived from fluorescence microscopy. These cells had ultrastructural features consistent with migratory neurons, like elongated nuclei, a centriole pair at the base of the leading process, and filopodia-like extensions at the end of the leading process.

This rich dataset revealed several unexpected phenomena. Migratory neurons were observed in a synapse-dense neuropil and made extensive contacts with various features of mature neurons including spines, axons, and dendrites. Migratory neurons also made frequent soma-soma contacts with mature neurons, which form large soma clusters in the avian striatum. Interactions between new neurons and mature neurons were asymmetrical, with migratory neurons deforming adjacent mature neurons and neuropil. In some instances, these indentations were so extreme that the new neurons appeared to tunnel through the clusters of mature neurons. These results reveal the value of applying EM connectomics to adult neurogenesis and suggest that migratory neurons may dramatically perturb the existing functional circuits as they migrate and integrate. Furthermore, they reveal the remarkable structural flexibility of mature neural circuits.

## Results

### Migratory neurons identified by morphological and ultrastructural features in EM dataset

To identify migratory neurons in the songbird basal ganglia connectome^23^ we created a library of ground truth morphologies of new neurons in the striatum using fluorescence microscopy in tissue sections (**Fig. S1)**. We used transgenic songbirds in which new neurons are sparsely labeled with GFP^25^, allowing visualization of the cellular morphology, combined with immunohistochemistry for doublecortin (DCX), a marker for migratory neurons^26,27^ or Hu to label neuronal identity^28,29^. These cells had two or more processes, and possessed a spindle-shaped soma with a thinner “trailing” process behind the soma, and a thicker leading process in front of the soma, which was often bifurcated (**Fig. S1,S2**). These cells resembled migrating neurons in other regions of the adult avian forebrain ^30^ and in the developing mammalian brain^31,32^.

We used a machine-learning based classifier, trained on a subset of manually identified cells, to identify other migratory neurons from the dataset (**Fig. 1a-d**; **Fig. S1;** see **Methods**) This approach defined a cell class consisting of 135 objects. Manual inspection revealed that 40 objects within this class met criteria for migrating neurons (false positives = 95/135; false negatives = 0; see **Methods**). We selected 35 true positives with largely complete reconstructions for further analysis (**Fig. S2**).

**Figure 1.**
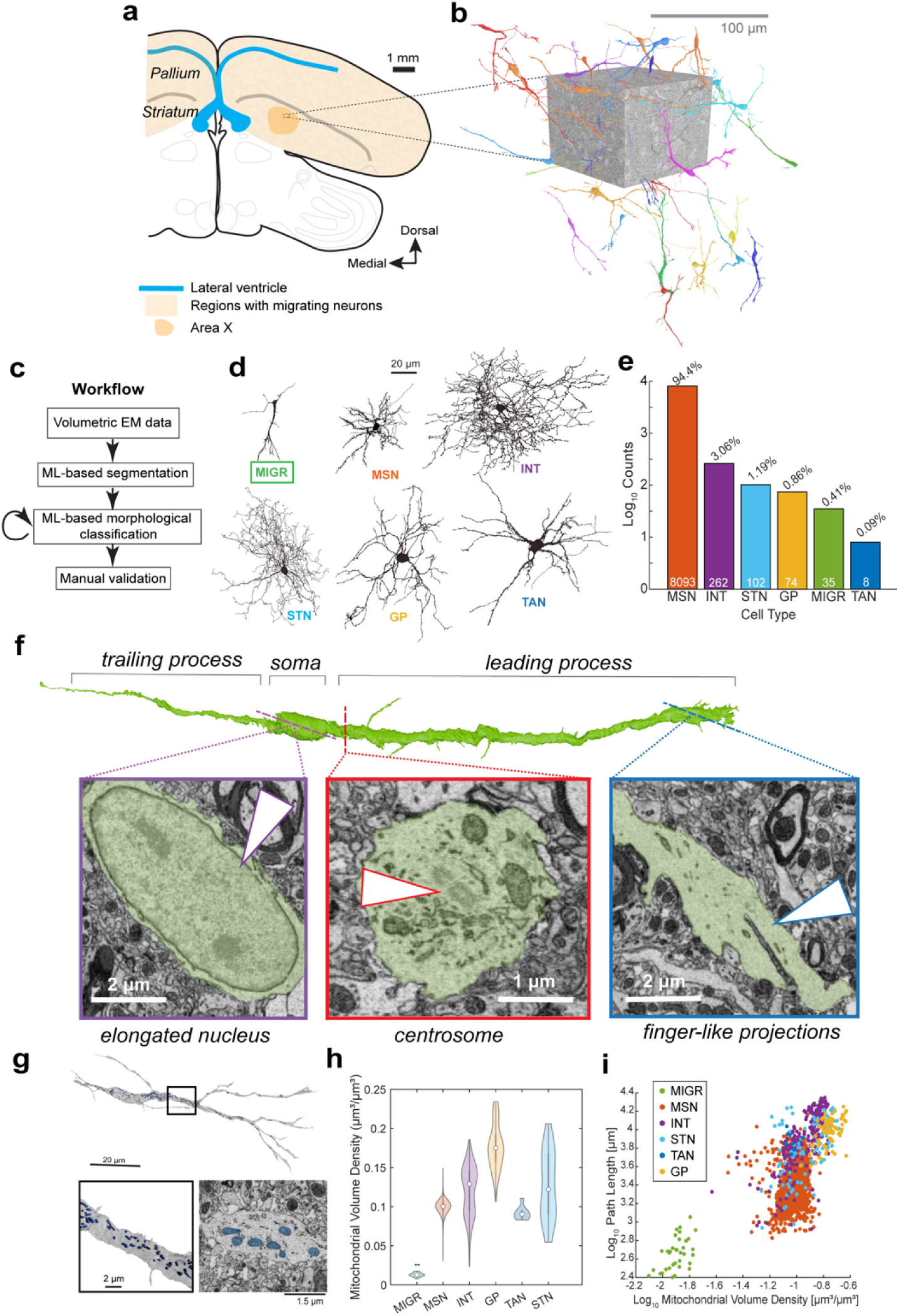
Identification of migratory neurons in volumetric EM. A) Overview of neuron migration in adult zebra finch brain. Neurons are born in the ventricular wall (blue line), and migrate throughout the pallium and striatum (orange shading), including Area X (orange outline). B) Reconstructions of migratory neurons from volumetric EM. C) Workflow for automated identification of migratory neurons. D) Tracings of representative cells from identified cell classes. E) Prevalence for each cell class in the data set. F) Ultrastructural features (arrows) characteristic of migratory neurons from an example cell. G) (top) Reconstruction of migratory cell with segmented mitochondria in blue. (bottom left) Zoom in on box region from top panel. (bottom right). Zoom-in cross section with mitochondria in blue. H) Mitochondrial densities across cell types. MIGR cell type has significantly lower mitochondrial densities than all other cell types (one-way ANOVA, p = 0). I) Path lengths and mitochondrial density volume for immature and mature neuron classes.

Identified cells had distinct morphological and ultrastructural features consistent with a migratory neuron identity (**Fig. 1f**). All cells had an elongated soma with thin cytoplasm surrounding a nucleus containing one or more nucleoli with heterochromatin-dense edges, as reported previously for migratory neurons across species and developmental stages ^29,33–37^. In 34/35 of these cells, the centrosome was clearly located below the nucleus at the base of the leading process, as expected for migratory neurons ^33,38,39^. A golgi apparatus was typically found near the centrosome, which has been reported to be aligned with the direction of neuron migration in immature neurons ^40–42^. All had one or more “finger-like” projections at the distal tips of their leading processes, similar to the filopodia or growth-cone structures described for migrating neurons and extending neurites ^43^. Finally, these cells typically had small, rounded mitochondria, which were clustered around the centrosome, as previously described for immature and migratory neurons ^44,45^ (**Fig. 1g**). They also exhibited a significantly lower mitochondrial density than mature neurons (**Fig. 1g-i**), perhaps indicative of a lower metabolic demand ^44,46–50^.

Together, our findings demonstrate that the songbird basal ganglia connectome^23^ contains a distinct class of cells with morphological and ultrastructural features that resemble immature neurons in the midst of or in the end stages of migration.

### Migratory neurons are present at a high density and oriented in multiple directions

Next, we evaluated the orientations and positions of identified migratory neurons throughout the connectome. We defined the orientation of the cell as a 3-dimensional vector beginning at the soma and ending at the first branch point of the leading process (**Fig. S3a,b; Methods**). Leading process vectors from different cells were oriented in a variety of directions (**Fig. S3a,d**). No directional bias was detected in the horizontal plane (Rayleigh’s test, p = 0.05) and a slight bias was detected in the azimuth plane (Rayleigh’s test, p < 0.001).

Identified migratory neurons were present at a density of 1,390 neurons per mm^3^ (35 cells in 0.025 mm^3^) and their distribution in the volume was well fit by a maximum entropy model, indicating uniform distribution throughout the tissue (**Fig. S3e**). Our findings of high soma position entropy and multidirectional leading process orientation are similar to observations from *in vivo* imaging studies of migration ^25,30^ and consistent with a diffusion-like process.

### Migratory cells are located within synapse-dense neuropil

Since migratory neurons were widely distributed throughout the tissue volume, we investigated whether they were located within regions rich in synapses, or, alternatively, constrained to low synapse density corridors (**Fig. 2a**). This connectome contains over 8 million high-confidence (>0.9 certainty) synapses at a density of 0.29 per µm^3^ (**Fig. 2b-d**). The local synapse density surrounding migratory neurons was not significantly different from the global synapse density (0.31 +/-0.07 per µm^3^, one-sample t-test, p = 0.12) or the density of synapses surrounding random points throughout the volume (0.28 +/-0.11 per um^3^, Kolmogorov-Smirnov test, p = 0.35, **Fig. 2e,f; Methods**), suggesting a roughly uniform probability distribution of synapses throughout the tissue. Indeed, the distribution of synapses throughout the entire volume was well fit by a maximum entropy model (uniform distribution, n = 8,278,615 simulated synapses) (**Fig. 2g-h**).

**Figure 2.**
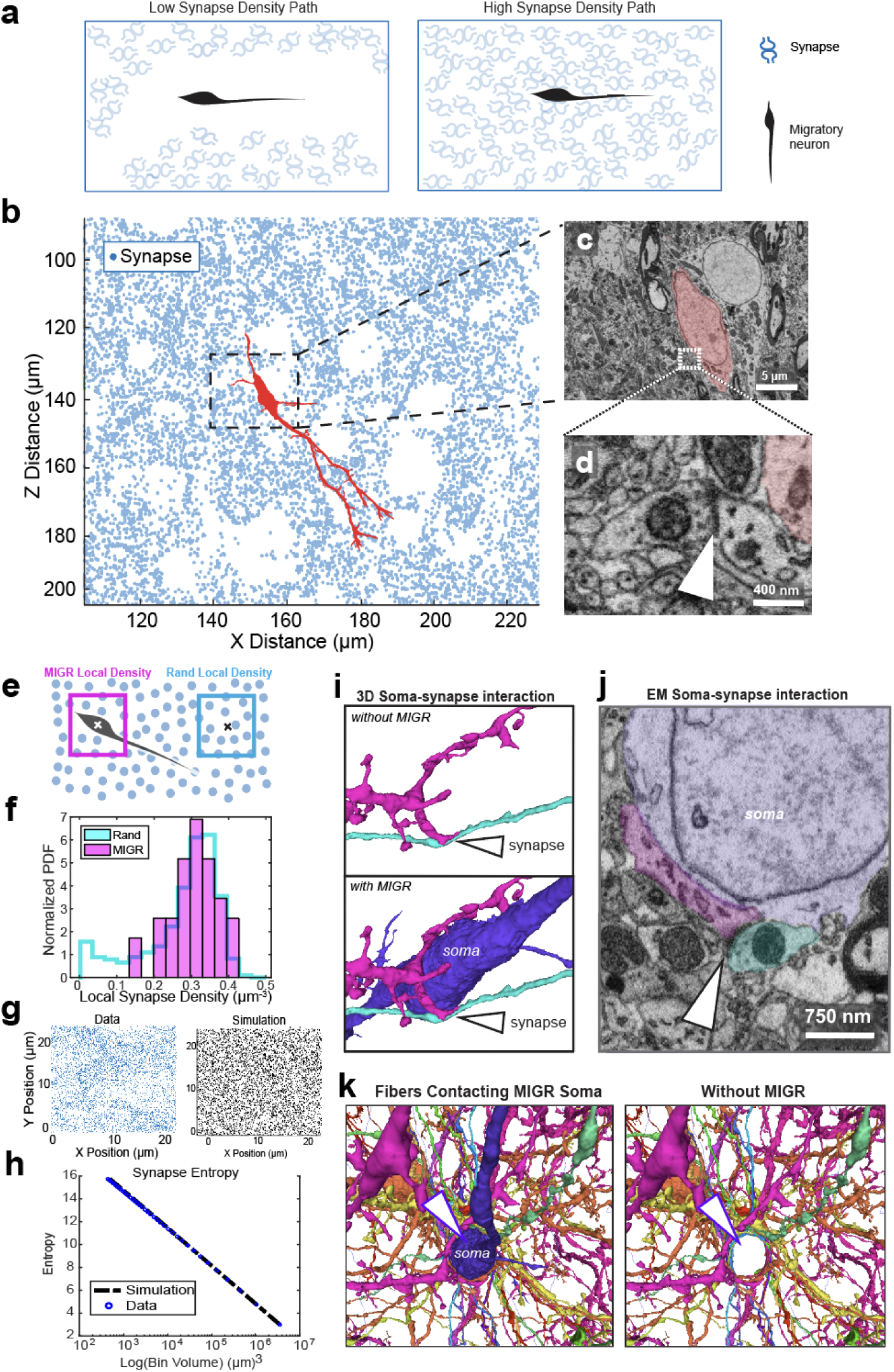
Migratory cells located within synapse-dense neuropil. A. Conceptual models: neurons may migrate through channels with low numbers of synapses (left) or through areas of high synapse density (right). B. Synapse locations (blue) surrounding a migratory neuron (MIGR) cell (red). C. Electron micrograph of dashed box in B. D. Zoom in on the dashed box region in C, showing synapse (arrow) near MIGR soma (red). E. Schematic for local density computation method. F. Histogram of local densities around MIGR cells (n = 35) and random points (n =29484). No significant difference was found between local MIGR density and local Rand density (K-S test, p = 0.35). G. (left) Synapse coordinate data in 25 um x 25 um x 10 um FOV (confidence >= 0.9; n = 4929 synapses). (right) Simulated synapse coordinates in 25 um x 25 um x 10 um FOV (n = 4929 simulated synapses). The simulation assumed a maximum entropy model and followed a uniform probability distribution. H. The Shannon Entropy of the synapse coordinates vs simulated uniform distribution across varying 3D bin sizes. I. (upper) 3D render of dendrite processes without MIGR cell. (bottom) 3D render of dendrite processes with MIGR cell. J. Zoom in on MIGR soma from I; two passing dendrites synapsing with each other (arrow) shown. K. EM micrograph of MIGR soma contacting synapse between two nearby dendrites (gray shaded box from J). L. (left) 3D render of MIGR soma making contact with several passing fibers. (right) same as left, but without MIGR visualization, showing concavity left in surrounding fibers.

Finally, visual inspection of ultrastructural features revealed that migratory neurons made contact with mature synapses (**Fig. 2i,j**). Moreover, we observed dendrites of mature neurons curving around the soma of migratory cells, raising the possibility that migratory neuron somata deform neuropil in their vicinity. Together, these findings suggest that new neurons are not preferentially migrating through low-synapse-density corridors, and instead migrate through neuropil highly dense in synapses.

### Migratory neurons interact with mature neuron somas

We next examined the interactions between new neurons and other classes of cells. In other systems, many new neurons use scaffolds, such as radial fibers or blood vessels to migrate ^34,51^. However, we observed only a small minority of migratory neurons (n = 8/35) associated with known migratory scaffolds (**Fig. S4**). In contrast, half of the identified migratory neurons formed close soma-soma associations with mature neurons (n = 17/35 migratory neurons; **Fig. S4**) and, to a lesser extent, astrocytes (n=4/35; **Fig, S5**). Interestingly, there was no significant difference between the proximity of migratory neurons to mature neurons (7.91 µm +/-2.55 µm) and the proximity of random points throughout the volume to mature neurons (9.14 µm +/-4.00 µm; KS test: p = 0.13; **Fig. S4i,j)**. Thus, these encounters may be due to the high density of mature neurons along the paths of migrating neurons, rather than an active process of attraction or avoidance.

### Migratory neurons tunnel through the mature avian striatum

Given the prevalence of mature-migratory neuron soma contacts, we examined their interactions in more detail. Specifically, we wondered whether migrating neurons conform their shape to the boundaries of mature neurons (**Fig. 3a**). Examination revealed that, rather than conforming to mature neurons, migrating neurons deformed adjacent neuron somas (n = 17 interactions; **Fig. 3b**).

**Figure 3.**
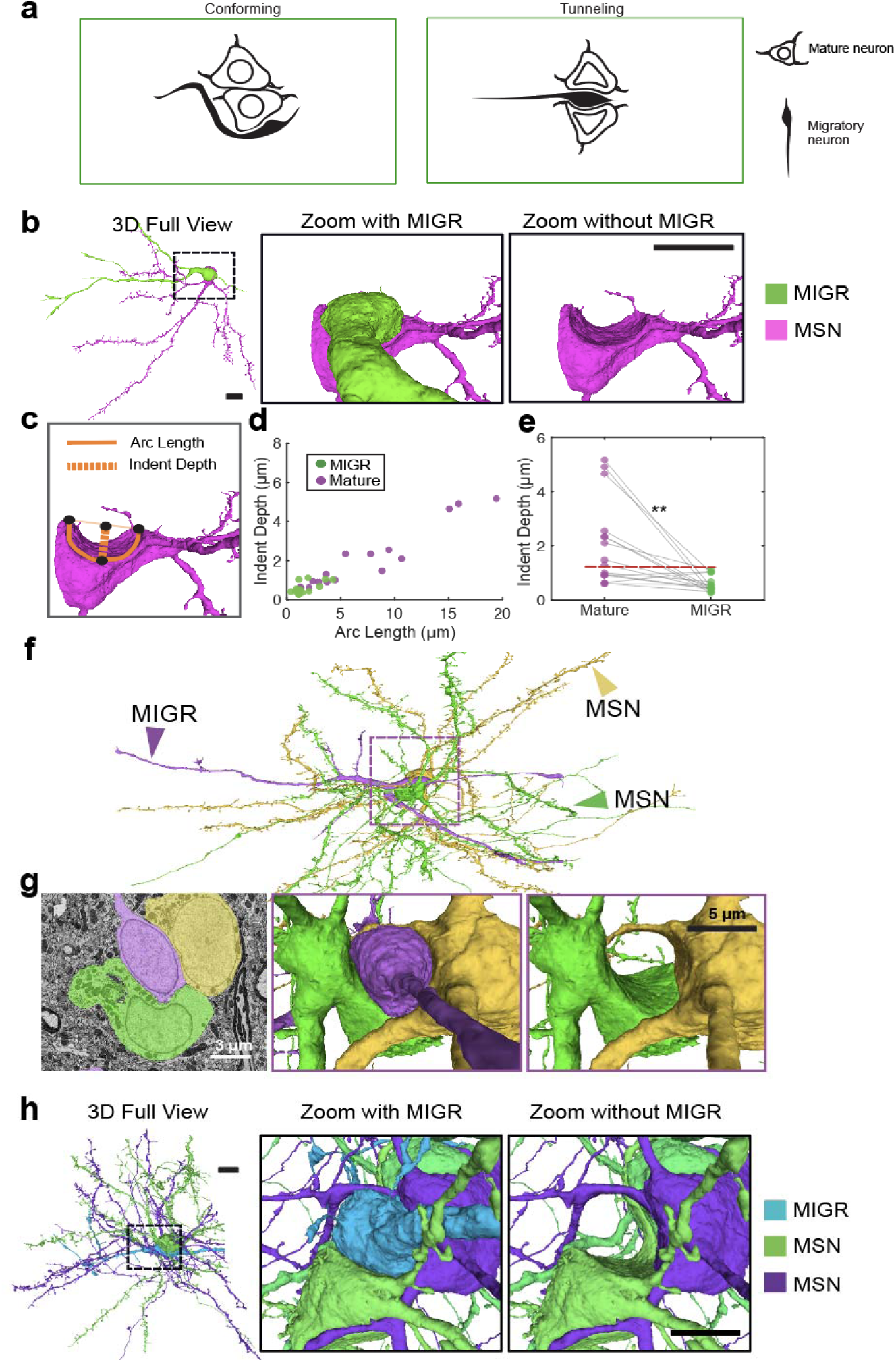
Migratory neurons form indentations in mature neuron somas. A) Alternative strategies for interactions with mature somas. B) (*left*) Example 3D render of soma-soma contact between a migratory neuron (MIGR; green) and one MSN (pink). (*middle*) zoomed in perspective of MIGR and MSN, and (*right*) same zoom in perspective without the MIGR cell. Depth of indent is 2.15 microns. Scale = 5 microns. C) Cell indent quantification method. D) Indent depth and length for each associated MIGR (green) and mature neuron (purple) (n = 17). E) Indent depth on each cell for MN-MIGR soma pairs (n = 17). Indent depth is significantly higher on mature neurons of each pair than their corresponding MIGR soma (paired t-test, p < 0.004). Red line indicates median mature neuron indent depth. F) Full 3D render of MIGR cell (purple) in soma-soma contact with two MSNs (yellow and green).G) (l*eft*) Electron micrograph of interaction from F. (*middle*) 3D render of MIGR cell (purple) and MSNs (green and yellow) from F. (*right*) 3D render of MSNs with MIGR cell removed. H) (*left*) “Tunneling” example with full view 3D rendering of MIGR (blue) and mature neurons (MSN class; green and purple), (*middle*) zoomed in perspective of MIGR and MSNs, and (*right*) same perspective as in middle without MIGR. Depth of indent is 4.91 microns. Scale = 5 microns.

To quantify this deformation we measured indentations at the site of contact between migrating and mature neurons **(Fig. 3c**). These indents were large and asymmetrical between contact partners (**Fig. 3d**). Indents were significantly deeper in mature neuron somas (2.03 +/-1.58 um deep, 7.15 +/-5.66 µm long) than in migratory neuron somas (0.59 +/-0.30 µm deep, 1.64 +/-1.08 µm long; paired t-test, p < 0.004; **Fig. 3e; Methods**). In several cases, migratory neurons were found within large concavities through one or more mature neurons. This phenomenon, in which new neurons appear to deform multiple adjacent neuron somas, axons, and dendrites, we refer to as “tunneling.” Tunneling was particularly evident in migratory neuron interactions with groups of densely packed mature neurons (**Fig. 3f-h**).

In the striatum, mature neurons were closer together than by chance (mean = 7.92 +/-2.78 µm; KS test, p < 0.00001) and frequently contacted the somas of other neurons (**Fig. 4**). Similar dense groupings of mature neurons have been reported previously in other areas of the avian brain where they have been called “clusters” ^8,52–55^. We found that 71.9% of all mature neurons (n = 6163) were found in clusters, defined as a connected graph of adjacent neuron somas (minimum distance <9 µm between soma centers, see **Methods**). Clusters (n=1574) ranged in size from 2 to 33 neurons (mean = 4 +/-3) and contained all classes of mature neurons (**Fig. 4e-h; Methods**) as well as glia (**Fig. S5**). The predominant cell type in the clusters were MSNs (n = 5786 MSNs; 93.9% of all clustered cells; **Fig. 4i**), however, after normalization by cell frequency, all neuron types had a similar contribution to cluster composition (**Fig. 4j**). New neurons were found in clusters, where they indented one or more mature neurons forming tunnels through the clusters (**Figure 4k-m**).

**Figure 4.**
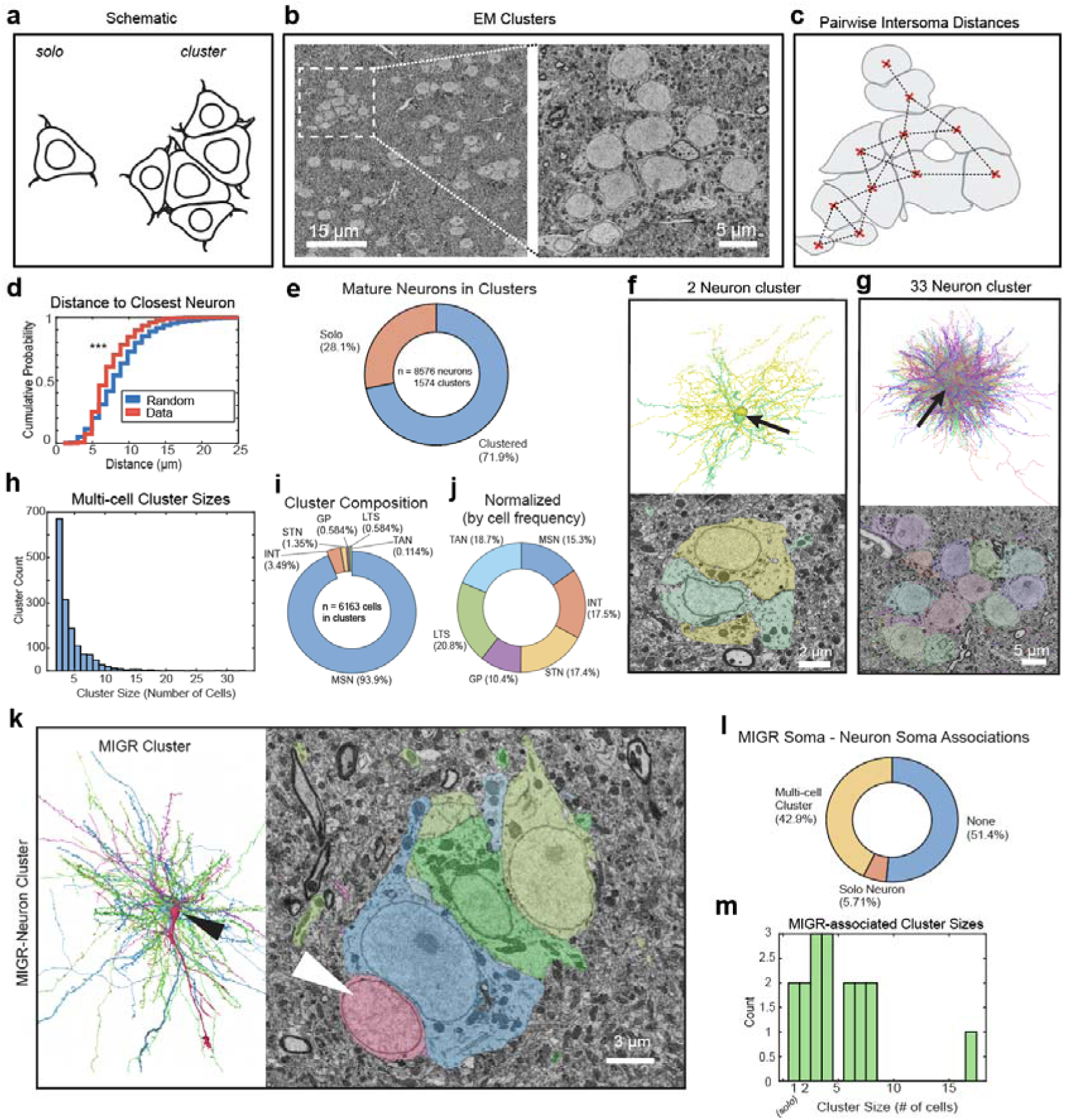
Migratory neurons associate with neuronal clusters. A) Schematic of solo neuron vs cluster of neurons. B) Single plane EM micrograph from Area X. (right) zoomed in view of the dashed region. C) Schematic for intersomal distance quantification. Clusters were identified using nearest neighbor network analysis based on intersomal distance. D) Mature neurons (blue line) are closer to neuron neighbors than expected by chance (red line; KS test, p < 0.00001). E) Of all mature neurons, 28.1% are on their own, and 71.9% are in clusters. F) Example two neuron cluster. 3D render (top, arrow) and EM Micrograph (bottom). G) Example 33 neuron cluster, 3d render (top, arrow) and EM plane through somas (bottom). H) Distribution of computed cluster sizes. I)Cell type composition of all clusters. J) Cell type composition of all clusters, normalized by the frequency of each cell type in the data. K) Example migratory neuron (MIGR) associating with cluster. 3D renders (left) and EM micrograph (right). Arrowheads point to MIGR soma. L) Frequency of associations of MIGR cells with mature neuron somas. M) Sizes of clusters contacting migratory neurons.

## Discussion

These data reveal the intricate interactions between migratory neurons in the adult striatum and their local environment. Our findings support a model in which migrating neurons disperse throughout dense neural tissue in multiple directions, making various contacts with surrounding structures. In addition, our data reveal a previously undescribed form of neuron migration in which new neurons cause deformities in nearby neurons and synapses. Together, these data highlight the value of applying connectomics to problems in developmental neuroscience ^56,57^.

Several features of the migration we observed resemble those reported in the embryonic and adult nervous system, including in humans ^35,58–62^. However, to our knowledge, tunneling migration by neurons has not previously been reported in the vertebrate nervous system. This behavior contrasts with chain migration in the rostral migratory stream ^63^, where neurons move through specialized corridors segregated from mature neurons, and with the amoeboid migratory strategy observed in some glial cells, which typically conform to the contours of mature neuronal somas ^64^. Interestingly, tunneling-like behavior has been described in metastatic cancer cells, which navigate confined spaces by actively deforming their microenvironments ^65^. Tunneling may therefore reflect a conserved strategy adopted by specialized migratory cell types in dense tissues. Future research could uncover the molecular and biophysical mechanisms that may enable this form of migration, and apply EM connectomics to evaluate its prevalence across systems.

Importantly, tunneling reveals a potential cost of adult neuron migration—disruption of existing circuits. As new neurons move through the mature tissues, mechanical disruption of cell bodies, axons and dendrites could affect synaptic connections, either through breakage or acute mechanical perturbation^66–68^. Such perturbations could alter circuit-level dynamics and may necessitate compensatory homeostatic mechanisms, similar to those proposed to support injury resilience in the songbird brain^69,70^. Alternatively, such network perturbations could also provide a substrate for plasticity. Indeed, recent advances in deep reinforcement learning indicate that sparse random perturbations in network connections can be used to facilitate continual learning in artificial neural systems^71,72^.

While the observed features are consistent with migratory identity, tunneling may reflect a terminal stage of migration and may lead to the formation of neuron clusters. In other areas of the songbird brain, new neurons preferentially integrate adjacent to mature ones ^8,30,52,73^, likely benefiting from some local cues that support survival, maturation and integration ^54,55,74^. Future studies using EM connectomics could illuminate these later stages, capturing how synaptic contacts evolve as new neurons integrate into functional adult circuits.

## Data availability

The dataset is available at SyConn Web (syconn.esc.mpcdf.mpg.de). All cells analyzed and used in figures can be visualized with SyConn Web using the Cell IDs given in **Table S1**.

## Ethics declarations

All authors declare no competing financial interests.

## Acknowledgements

We thank Michale Fee, JoAnn Buchanan, Forrest Collman, and Maya Medalla for helpful discussions. We thank Shawn Sorrells for comments on the manuscript. We thank the Boston University Neurophotonics Center for funding, technical support, and imaging facilities. We also acknowledge the Boston University Animal Science Center Staff for their care and maintenance of the zebra finch colony, and we give a special acknowledgement to the sacrifice of the zebra finches included in this work.

## Author contributions

JK performed data collection, data curation and managed data sharing. BBS and JK conceived the project. NRS, AR, SJC and DS performed analysis and contributed to figures. NRS wrote the manuscript with feedback from BBS. All authors discussed and edited the manuscript.

## Supplementary Materials for

**Includes:**

Methods

Figs. S1 to S6

Table S1

Supplementary References

## Methods

### Immunohistochemistry

Animal use procedures were approved by the Boston University Institutional Animal Care and Use Committee (IACUC; Protocol #201800577) and carried out in accordance with National Institutes of Health standards. Birds were housed under standard conditions (12h/12h light/dark cycle).

Transgenic UBC-GFP transgenics^1,2^ were bred at Boston University. Adult male zebra finches were euthanized with Euthasol (Virbac) and perfused with up to 20 mL of 1X phosphate-buffered saline (PBS), followed by up to 50 mL of 10% neutral buffered formalin. The brain was extracted and stored in formalin overnight. After fixation, the brain was washed with PBS and sliced into 40 or 50 μm sections with a vibratome (Leica; VT1000S).

The primary antibodies used were a mouse monoclonal anti-DCX (1:300 dilution) (Santa Cruz Biotechnology, sc-271390 (E−6), lot: C3121) ^3^. We also stained with a mouse monoclonal anti-HuC/HuD that is a marker for cells in neuronal lineage in 1:250 dilution (ThermoFisher Scientific A-21271)^4^. The secondary antibodies used were Alexa Fluor 568 conjugated goat anti-mouse (Invitrogen) in a 1:500 dilution.

Antigen retrieval was performed prior to staining by incubating slices in a 50 mM solution of Tris buffer (Fisher Scientific) at a pH of 8.4 for 30 min. Then, tissue sections were placed in a blocking solution composed of 2% nonfat dry milk, 0.02% Triton X-100 (Sigma), and 1X PBS. Sections were incubated at 4°C overnight in primary antibody diluted in blocking solution, washed 3 times with PBS, then incubated with secondary antibody diluted in blocking solution for 1 h at room temperature or overnight at 4°C. Sections were mounted on microscope slides (Fisher Scientific) using Vectashield Antifade Mounting Medium with DAPI (Vector Laboratories, Inc.). Histological fluorescent images from striatal nucleus Area X were acquired using a confocal microscope (Zeiss LSM800, Germany). Image processing was performed with FIJI/ImageJ.

### Machine learning based identification of migratory neurons

Ground truth cells (n=14) were manually selected from the zebra finch Area X connectome^5^ using morphological characteristics shared with fluorescent imaging data identifying migratory neuron morphology. These cells were used to train a supervised learning algorithm, based on previous methods^6–8^. The IDs for these cells were: 751594441, 971410555, 2157811112, 38332384, 1773210639, 328158994, 1644151292, 561503453, 1104854425, 28511657, 2656656549, 669941616, 763200873, 245868794.

This approach returned 135 objects which were manually inspected. 40/135 exhibited morphological and ultrastructural features consistent with migratory neurons. This data indicate a false positive rate of 79.4%.

To evaluate the false negative rate, i.e. migratory neurons that were missed by the algorithm, we visually inspected a set of >2,000 low-confidence (< 0.5 certainty) medium spiny neurons and microglia, possessing a segment volume between 460 and 1500 µm^3^, and < 200 identified synapses. Using this method we identified 28 cells with morphology and ultrastructure features consistent with migratory neurons. All 28 cells found through a manual algorithm were found within these 40, indicating an undetectable false negative rate. If cells were drastically clipped, or had no distinct leading process (5/40), they were excluded from further analysis.

### Cell orientation analysis

To determine the orientation vectors of each cell, three X, Y, Z coordinates were extracted in the Syconn Neuroglancer interface. The three coordinates corresponded to the start of the soma (*a_XYZ_*; at the base of the leading process), the first bifurcation point of the leading process (*b_XYZ_*), and the distal tip of the leading process (*c_XYZ_*; **Fig. S3a,b**). All further analyses were performed in Matlab using a custom script.

The orientation of the cell was determined by the direction of the vector between *a_XYZ_* and *b_XYZ_* and the magnitude of the vector (only used for plotting in **Fig. S3c,d**) was determined by the approximate length of the cell from *a_XYZ_* to *c_XYZ_*.

After the orientation vector was extracted (*b_XYZ_* – *a_XYZ_*), it was normalized by dividing the vector by its Euclidean norm (using the *norm* Matlab function) to make the value of each dimension (in X, Y, and Z) its relative contribution to the overall orientation (maximum value of 1 in any dimension). If all cells were pointed in the same direction, the magnitude of their averaged orientation vector would be 1. This was to account for differences in physical length between the soma and bifurcation point of all cells which should not contribute to the overall orientation estimates.

### Mitochondrial density estimates

Mitochondria were segmented and annotated previously ^7^. For cell IDs that were filtered and verified above, the summed volume of all mitochondria voxels was divided by each segmented cell’s volume to yield the total mitochondrial volume density (µm^3^/µm^3^). A one-way ANOVA was performed to test for significant differences in mitochondrial density across groups.

### Synapse density estimates

Synapses were segmented and annotated previously using automated methods ^7,8^. Only synapses with a classifier certainty greater than or equal to 0.90 were used for analysis in Matlab. For local density estimates around migratory neurons (MIGR), all synapses within a 10 µm search radius from the MIGR soma center coordinate (n = 35 MIGR somas) were counted and divided by the search volume (20 µm x 20 µm x 20 µm) to yield the local synapse density in µm^-3^ for each cell. For local synapse density around random points, the local synapse density was computed around every micron voxel throughout the data (n = 29484).

### Shannon Entropy estimation

Shannon entropy of actual data (soma and synapse positions) or simulated data was computed as described previously^1^. Briefly, the three-dimensional volume containing the positions was binned into different sized bins, and we computed the probability of containing a data point for each 3-dimensional bin. We then input these probabilities into the Shannon entropy formula:

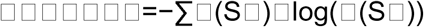

where p(S□) is the probability that a soma’s or synapse’s position is within any spatial bin. We repeated this process while we increased the bin sizes and plotted the entropy across bin sizes.

### Cell somas and cluster analysis

Soma positions were extracted from segmented objects with several predicted compartments. Only cells that had a soma compartment, as well as dendrites and an axon that were at least 200 µm in skeleton path length were selected. All cell types other than MSN and INT were then manually verified and any segmented cells that were merged with another fragment or cell were removed from analysis. For MSN and INT, a random subset of cells was verified manually. The median of the soma vertex coordinates was used as the “soma center.” Three-dimensional coordinates from each segmented soma were then analyzed in Matlab (n = 8576 neurons).

To find the nearest neighbor of all mature neurons, pairwise distance calculations were performed between each soma center coordinate. The minimum pairwise distance for each cell was considered the distance to its nearest neighbor. A similar analysis was performed for MIGR cells for which the cell soma center was determined manually in the SyConn Neuroglancer interface. Then, the minimum pairwise distance between MIGR and all other mature neurons was extracted to find the distance to the nearest neighbor. The minimum distance found between a MIGR soma center and a mature neuron soma center was 2.575 µm.

For “random” distances, we created a binary three-dimensional matrix where the position of each soma center coordinate was a “1”. Then, we performed a distance transform on this binary 3D soma matrix to find distance between every element and the nearest nonzero element. Distances below 2.575 were determined to be biologically improbable and were removed.

For an analysis of cluster size and composition, a graph-based connected components approach was taken. First, a distance transform on the binary 3D matrix of soma positions was performed. The resultant matrix contained the 3D distance between each element and every other nonzero element. We next iteratively determined an appropriate distance threshold, 9 µm (approximately double the average soma radius of 4.46 µm) to find cells belonging to the same cluster. After testing several distance thresholds ranging from 8 to 11 µm, we found that 9 µm yielded no false positives in the largest output clusters. However, using this approach, we detected a nonzero false negative rate, suggesting we are underrepresenting cluster sizes with this analysis.

Then, we applied this threshold to the distance matrix so that every value greater than 9 µm became zero, and cells that were clustered together below this distance threshold were represented as groups of ones. Using Matlab’s conncomp function, connected components composed of singular groups of ones were found and stored in structs as separate clusters. The cell IDs belonging to these clusters were extracted and visualized in the Syconn Neuroglancer interface to confirm accurate cluster identification with this algorithm. Cluster sizes were computed from the data and cluster compositions were extracted.

**Figure S1.**
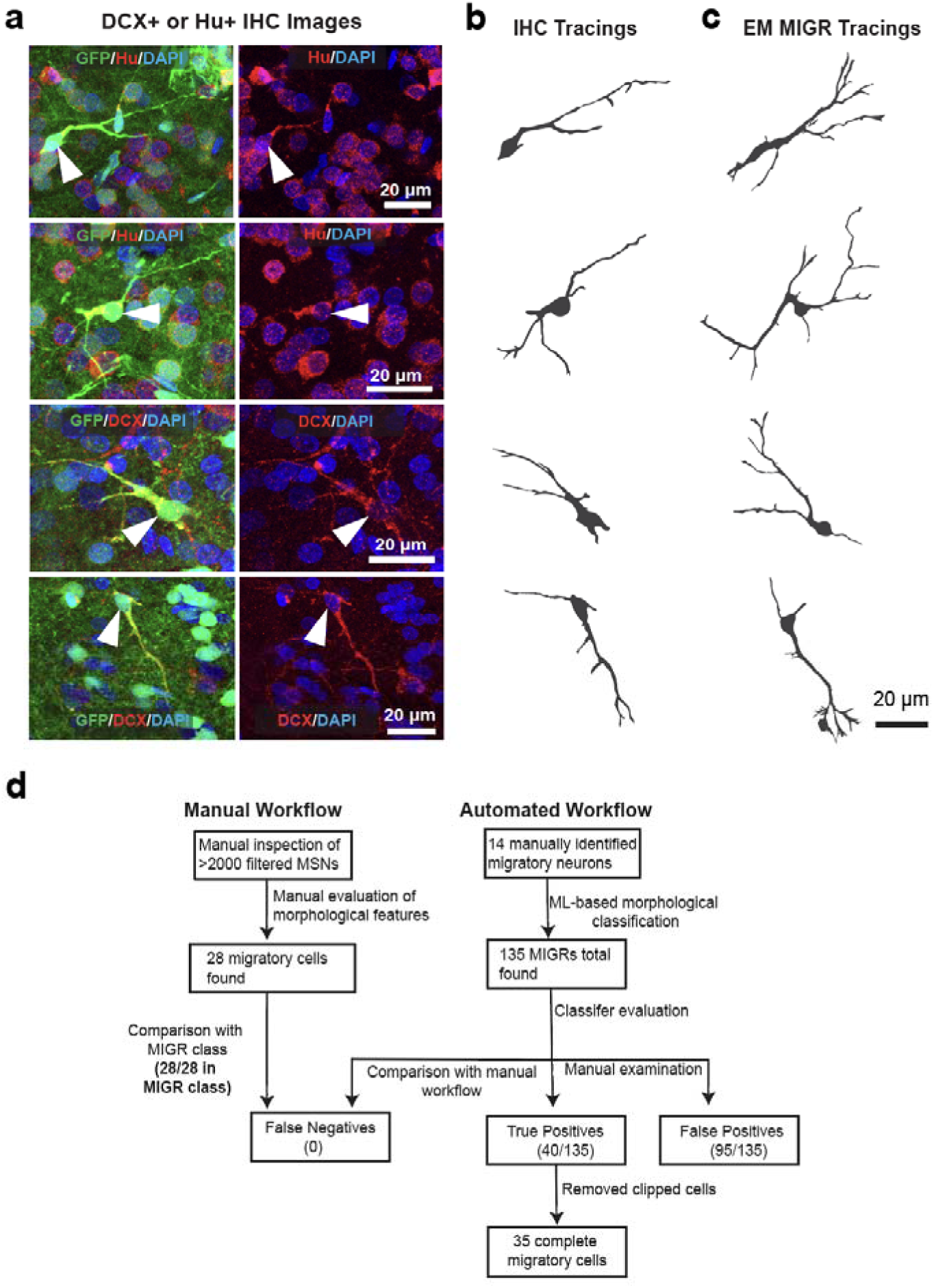
Workflow for identification of migratory neurons in EM. A) Confocal images of histological sections from UBC-GFP transgenic birds. Sections are labeled with immunohistochemistry against GFP (green) and either DCX (red, upper two panels) or HU (red, lower panels). Co-stained cells with a migratory morphology are indicated with an white arrow. B) Tracings of cells from A. C) Reconstructions of cells with similar morphology from the songbird connectome. D) Two workflows were used to identify migratory cells in the songbird connectome: manual (left) and automated (right). In the manual workflow, expert human observers inspected hundreds of reconstructions to identify cells that met the morphological criteria of migrating neurons. In the automated workflow, 14 manually identified cells were used to train a convolutional neural network-based classifier. The trained classifier resulted in the construction of a cell class with 135 objects. The manual workflow was then used to assess the false negative rate of the automated approach.

**Figure S2.**
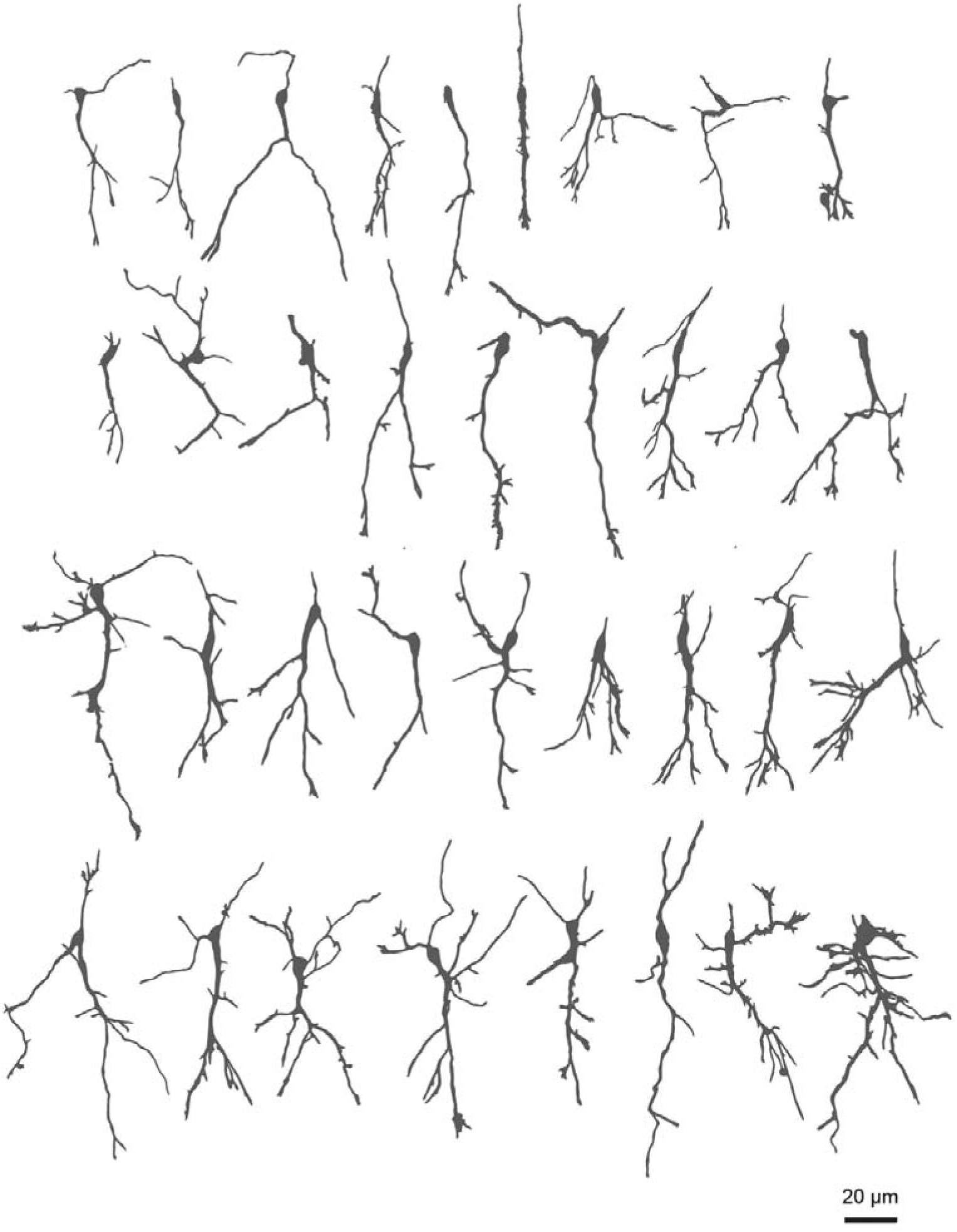
Morphology of identified migratory cells. Cells with migratory neuron morphology and ultrastructure ordered by volume. Note cells vary in complexity; most possess branched leading processes.

**Figure S3.**
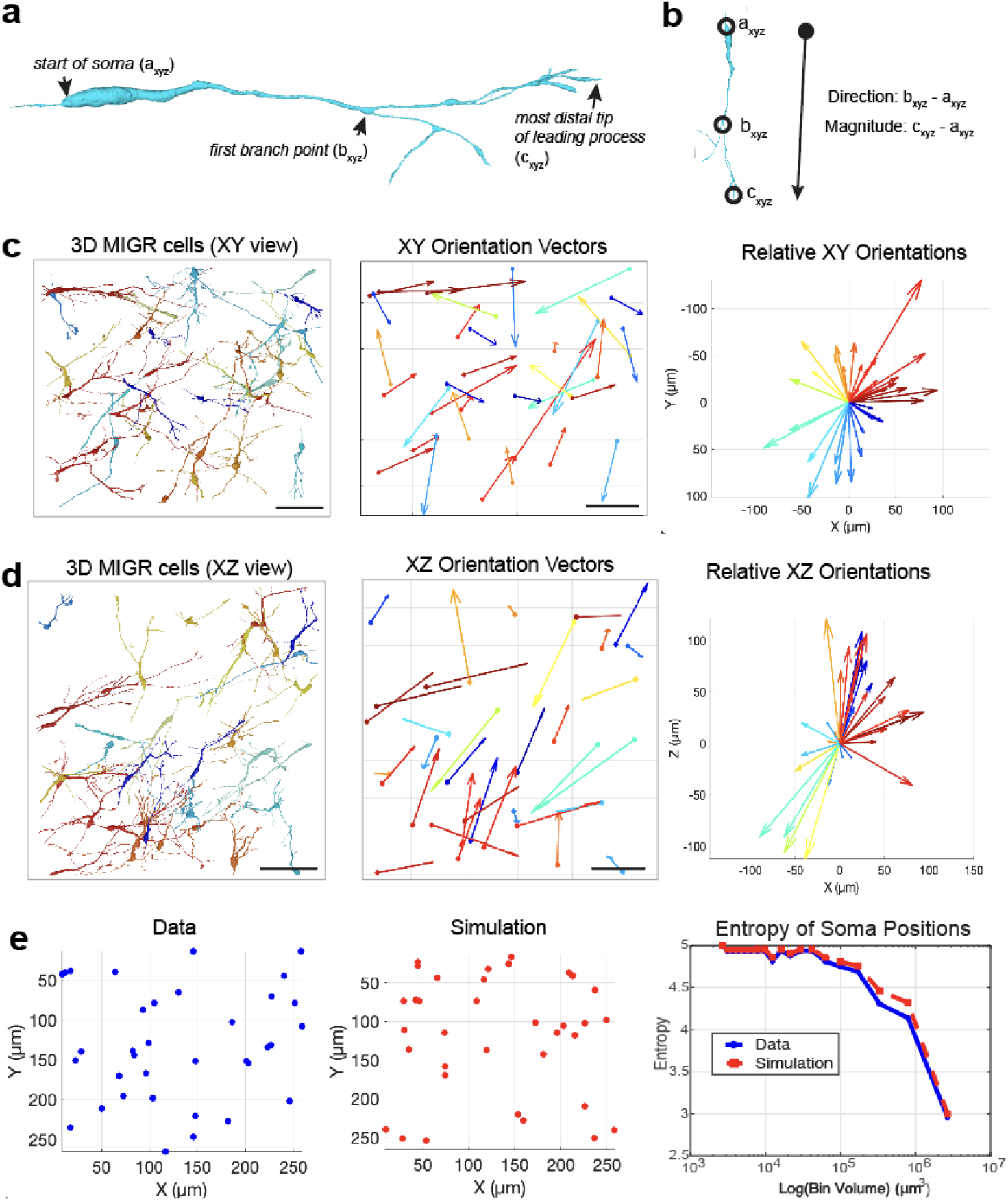
Migratory neurons are widely distributed and oriented in multiple directions. **A**. Morphological features used for computing cell orientation. **B**. Formula for computing cell orientation. **C**. Left: XY view of migratory neurons (MIGR; n=35) in the volume. Middle: XY orientation vectors; magnitude of vectors vary according to length of MIGR. Scale bars: 50 microns. Right: Polar plot of relative XY vector directions with start position and magnitude normalized. No directional bias was detected (Rayleigh’s test, p = 0.05). **D**. Left: XZ view of MIGR cells (n=35) i. Middle: MIGR XZ orientation vectors; Right: Polar plot of relative XZ vector directions. Significant directional bias was detected (Rayleigh’s test, p < 0.001). **E**. Left: Recorded soma positions. Middle: Simulated soma positions (maximum entropy model). Right: Computed entropy of data (blue) vs simulation (red) across different bin sizes.

**Figure S4.**
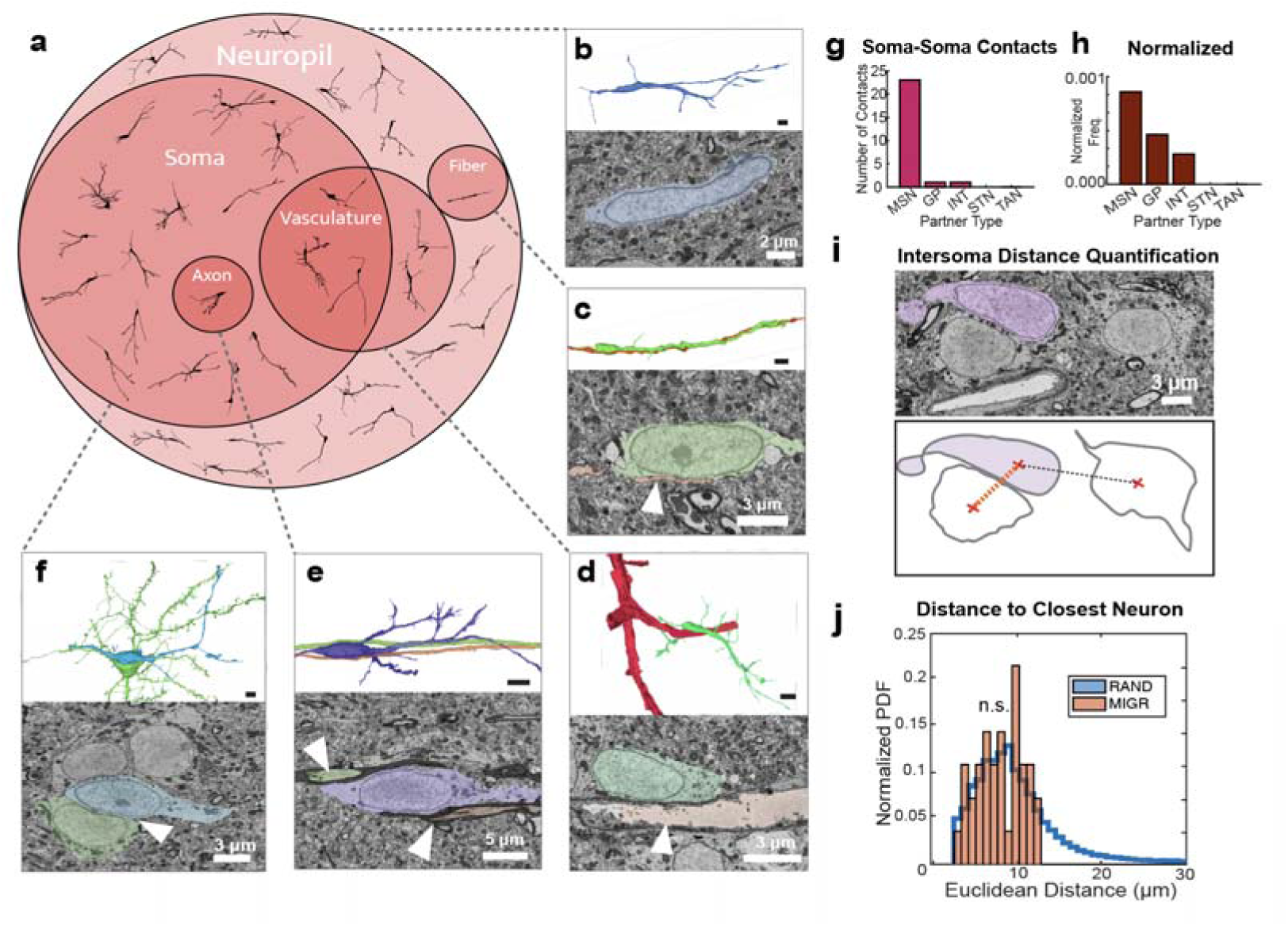
Migratory cells contact a variety of mature cell types. a) Partners for migratory cells: We evaluated the contacts of all migratory neurons to test if they associate with a particular type of structure known to be a cellular scaffold for migration. We manually inspected the microenvironment surrounding the somas of migratory neurons in three dimensions and recorded adjacent cells and structures with significant membrane contact as determined by manual inspection.. Circle size represents the number of migratory cells (shown in black) that make contact with a particular structure. Overlapping circles indicate cells with multiple contact partners for cells in the intersecting region. b) Example migratory neuron (MIGR) found in neuropil with no observable soma-soma contacts nor fibers closely aligned with the cell soma. c-f) Example cells with contact partners shown as a 3D rendering (top) and color-coded 2D electron micrograph (bottom). Contact partners include fibers [c], vasculature (f), axons (e), mature neurons somas (f) g) Frequency of MIGR soma contacts with other neuron types. h) Normalized frequency of MIGR soma contacts with multiple neuron types. i) Method of quantification of intersoma distances between MIGR cells and mature neurons. Red X’s denote the 3D center coordinate for each soma, and the dashed lines represent 3D distance. Orange dashed line is the minimum intersoma distance. j) Computed distances of random points (RAND; blue) to nearest mature soma center, and distances of 35 MIGR soma centers to nearest mature soma center (MIGR; orange). No significant difference in distribution (KS test, p = 0.15).

**Fig. S5.**
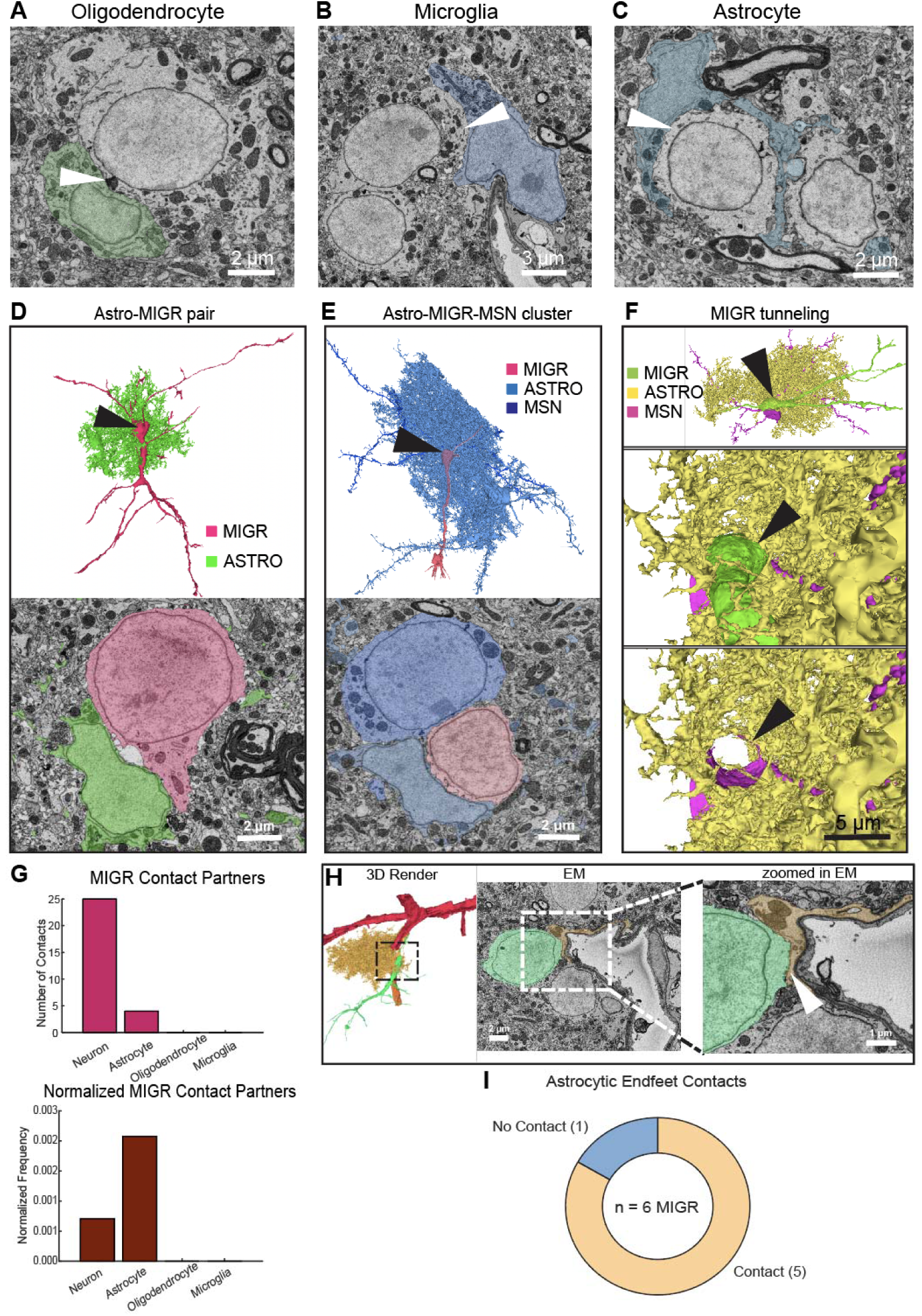
Glial interactions with mature and migratory neurons. A-C) Example soma-soma interaction between oligodrendrocytes (A), microglia (B) and Astrocytes (C) and medium spiny neurons (MSNs). Arrowhead points to the deformation of the glia soma membrane. D) Soma-soma interaction between migratory neuron (MIGR) and astrocyte. 3D render (top) with an arrow pointing to somas portrayed in EM micrograph (bottom). E) MIGR cell part of a soma cluster containing an MSN and an astrocyte. 3D render (top) with an arrow pointing to somas portrayed in EM micrograph (bottom). F) Example of MIGR tunnel through MSN and astrocyte cluster. Top panel shows zoom out 3D render with arrowhead pointing to MIGR soma. Middle panel shows zoom in view of soma interaction between all cells. Bottom panel shows same view as middle without the MIGR cell. G) (top) Bar plot of MIGR soma contacts with neurons and different glial types. (bottom) Normalized by cell frequency. H) (left) 3D render of MIGR cell contacting portion of blood vessel. (middle) EM micrograph of MIGR soma contact with blood vessel. Zoom in on dashed box from middle. White arrowhead points to astrocytic endfeet between MIGR soma and endothelial cells of vessel. I) Proportion of MIGRs that contact astrocytic endfeet between MIGR and vessel.

**Figure S6.**
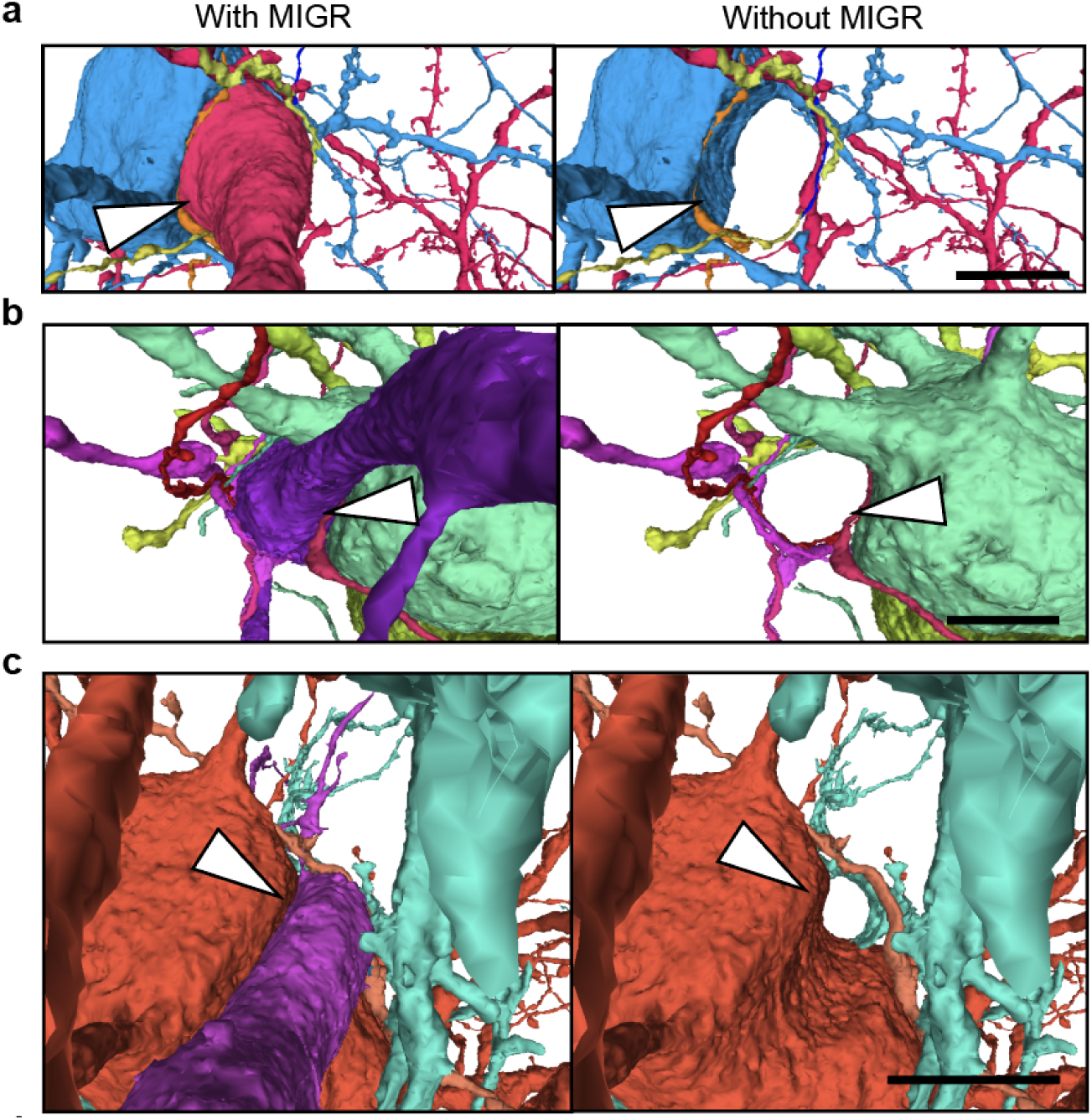
Tunneling examples. Each row is a zoom in on migratory neuron (MIGR) soma interaction with surrounding neurons or neuropil. Left column has MIGR rendered, right column is the same view shown without the MIGR cell. White arrowhead points to MIGR soma position. MIGR IDs from panels a respectively, are 1977673147, 299547481, 1711566114, and 2732540650. Scale is 5 µm.

**Table S1.**
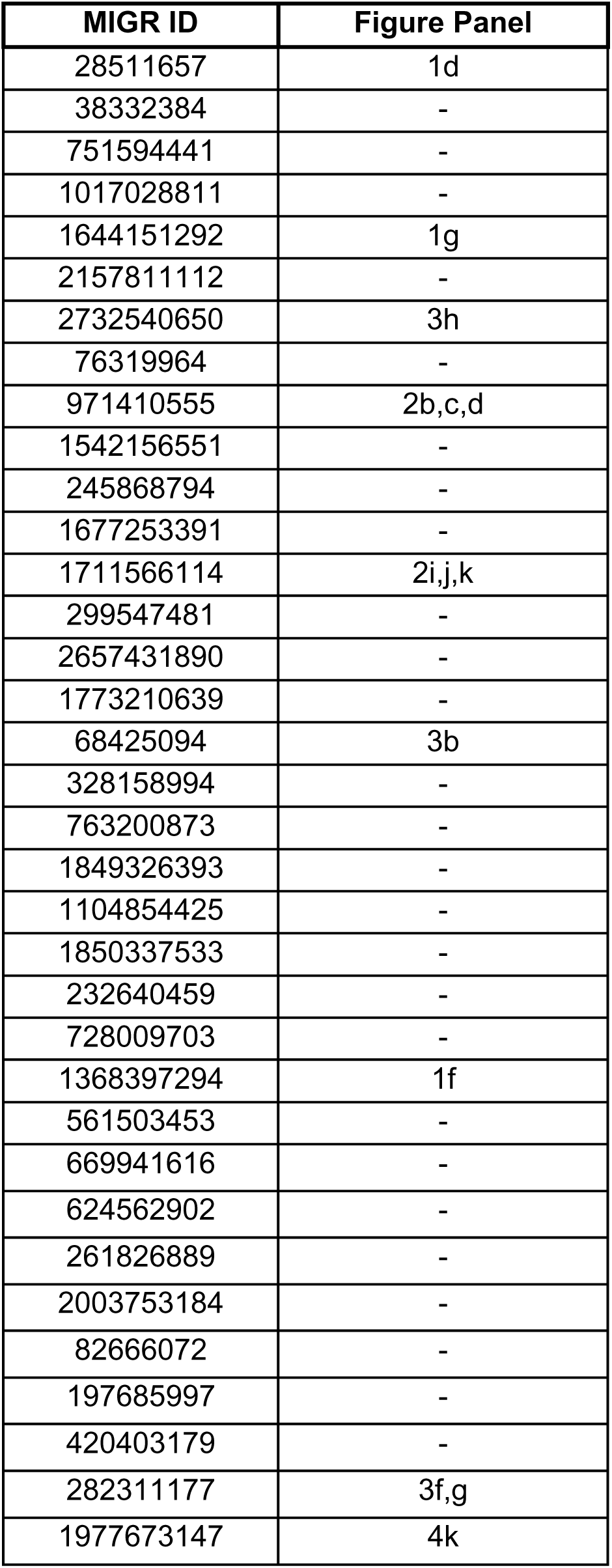
Table of identified migratory neurons. ID numbers of the 35 complete migratory cells (MIGR) and associated main figure panel. All of the cells are depicted in Fig. 1b.

